# Respiratory Alkalosis Provokes Spike-Wave Discharges in Seizure-Prone Rats

**DOI:** 10.1101/2021.08.15.456408

**Authors:** Kathryn A. Salvati, George M.P.R. Souza, Adam C. Lu, Matthew L. Ritger, Patrice G. Guyenet, Stephen B. Abbott, Mark P. Beenhakker

## Abstract

Hyperventilation reliably provokes seizures in patients diagnosed with absence epilepsy. Despite this predictable patient response, the mechanisms that enable hyperventilation to powerfully activate absence seizure-generating circuits remain entirely unknown. Using the WAG/Rij rat, an established rodent model of absence epilepsy, we demonstrate that absence seizures are highly sensitive to arterial carbon dioxide, suggesting that seizure-generating circuits are sensitive to pH. Moreover, hyperventilation consistently activated neurons within the intralaminar nuclei of the thalamus, a structure implicated in seizure generation. We show that intralaminar thalamus also contains pH-sensitive neurons. Collectively, these observations suggest that hyperventilation activates pH-sensitive neurons of the intralaminar nuclei to provoke absence seizures.

## Introduction

Epilepsy is a common neurological disorder characterized by recurrent and spontaneous seizures. Yet, accumulating evidence indicates that seizures are not necessarily unpredictable events (Amengual-Gual et al., 2019; Bartolini & Sander, 2019; Baud et al., 2018; Ferlisi & Shorvon, 2014). Several factors affect seizure occurrence, including metabolism (Lusardi et al., 2015; Masino et al., 2012; Masino & Rho, 2012, 2019), sleep (Bazil, 2019; Fountain et al., 1998; Malow et al., 1999; Nobili et al., 2001), catamenia (Herzog & Frye, 2014; Joshi & Kapur, 2019; Reddy et al., 2001), light (Padmanaban et al., 2019) and circadian rhythm (Amengual-Gual et al., 2019; Debski et al., 2020; Smyk & van Luijtelaar, 2020; Stirling et al., 2021). In extreme cases, stimuli immediately provoke seizures, a condition known as *reflex epilepsy* (Kasteleijn-Nolst Trenité, 2012; Koepp et al., 2016). The mechanisms that render certain seizure-generating networks susceptible to external factors remain unknown.

A highly reliable seizure trigger associated with childhood absence epilepsy is hyperventilation. Between 87-100% of all children diagnosed with the common *Genetic Generalized Epilepsy* produce spike-wave seizures upon voluntary hyperventilation (Hughes, 2009; Ma et al., 2011; Sadleir et al., 2009). Indeed, hyperventilation serves as a powerful tool for diagnosing this childhood epilepsy (Adams & Lueders, 1981; Holowach et al., 1962; Sadleir et al., 2006; Watemberg et al., 2015). Remarkably, as no single genetic etiology drives absence epilepsy (Chen et al., 2013; Crunelli & Leresche, 2002; Helbig, 2015; Koeleman, 2018; Robinson et al., 2002; Xie et al., 2019), hyperventilation appears to recruit fundamental seizure-generating mechanisms shared by virtually all patients.

Exhalation of CO_2_ during hyperventilation causes hypocapnia, a state of decreased arterial CO_2_ partial pressure (PaCO_2_), and respiratory alkalosis, a state of elevated arterial pH (Laffey & Kavanagh, 2002). Hyperventilation also causes rapid arterial vasoconstriction (Raichle & Plum, 1972) and increased cardiac output (Donevan et al., 1962). Recent work demonstrates that inspiration of 5% CO_2_ blunts hyperventilation-provoked spike-wave seizures in humans (Yang et al., 2014). Collectively, these observations suggest that respiratory alkalosis serves as the primary trigger for hyperventilation-provoked absence seizures.

Spike-wave seizures associated with absence epilepsy arise from hypersynchronous neural activity patterns within interconnected circuits between the thalamus and the cortex (Avoli, 2012; Beenhakker & Huguenard, 2009; Huguenard & McCormick, 2007; McCafferty et al., 2018; McCormick & Contreras, 2001; Meeren et al., 2002). The crux of the prevailing model describing absence seizure generation includes an initiating bout of synchronous activity within the somatosensory cortex that recruits rhythmically active circuits in the thalamus (Meeren et al., 2002; Sarrigiannis et al., 2018). With widespread connectivity to the cortex, the thalamus then rapidly generalizes spike-wave seizures to other brain structures. The extent to which thalamocortical circuits respond to shifts in pH during hyperventilation-induced respiratory alkalosis is unknown.

Herein, we test the hypothesis that respiratory alkalosis regulates the occurrence of spike-wave seizures. We demonstrate that hyperventilation-provoked absence seizures observed in humans can be mimicked in an established rodent model, the WAG/Rij rat (Coenen, 2003; Coenen et al., 1992; Russo et al., 2016; van Luijtelaar & Coenen, 1986). We first show that hyperventilation induced with hypoxia reliably evokes respiratory alkalosis and increases spike-wave seizure count in the WAG/Rij rat. When supplemented with 5% CO_2_ to offset respiratory alkalosis, hypoxia did not increase spike-wave seizure count. Moreover, hypercapnia alone (high PaCO_2_) reduced spike-wave seizure count despite a robust increase in respiration rate. We also show that optogenetic stimulation of brainstem respiratory centers to produce respiratory alkalosis during normoxia induces CO_2_-sensitive spike-wave seizures. Collectively, these results identify respiratory alkalosis as the primary seizure trigger in absence epilepsy following hyperventilation. Finally, we show that structures of the intralaminar thalamic nuclei are both (1) activated during respiratory alkalosis, and (2) pH-sensitive. Thus, our data demonstrate that respiratory alkalosis provokes spike-wave seizures and shine a spotlight on the poorly understood intralaminar thalamus in the pathophysiology of spike-wave seizures.

## Results

### Hypoxia triggers spike-wave seizures in the WAG/Rij rat

We first set out to determine if an accepted rat model of absence epilepsy, the WAG/Rij rat, recapitulates hyperventilation-provoked absence seizures, as observed in humans. We combined whole-body plethysmography and electrocorticography/electromyography (ECoG/EMG) recordings in awake WAG/Rij rats to assess respiration and spike-wave seizure occurrence while exposing animals to different gas mixtures of O_2_, CO_2_ and N_2_ (Figure 1A,B). We only considered spike-wave seizures that persisted for a minimum of two seconds and occurred concomitantly with behavioral arrest in the animal. Spike-wave seizures are distinguishable from non-REM sleep based on the appearance of 5-8 Hz frequency harmonics in the power spectrogram (see Figure 1B, *expanded trace*).

**Figure 1.**
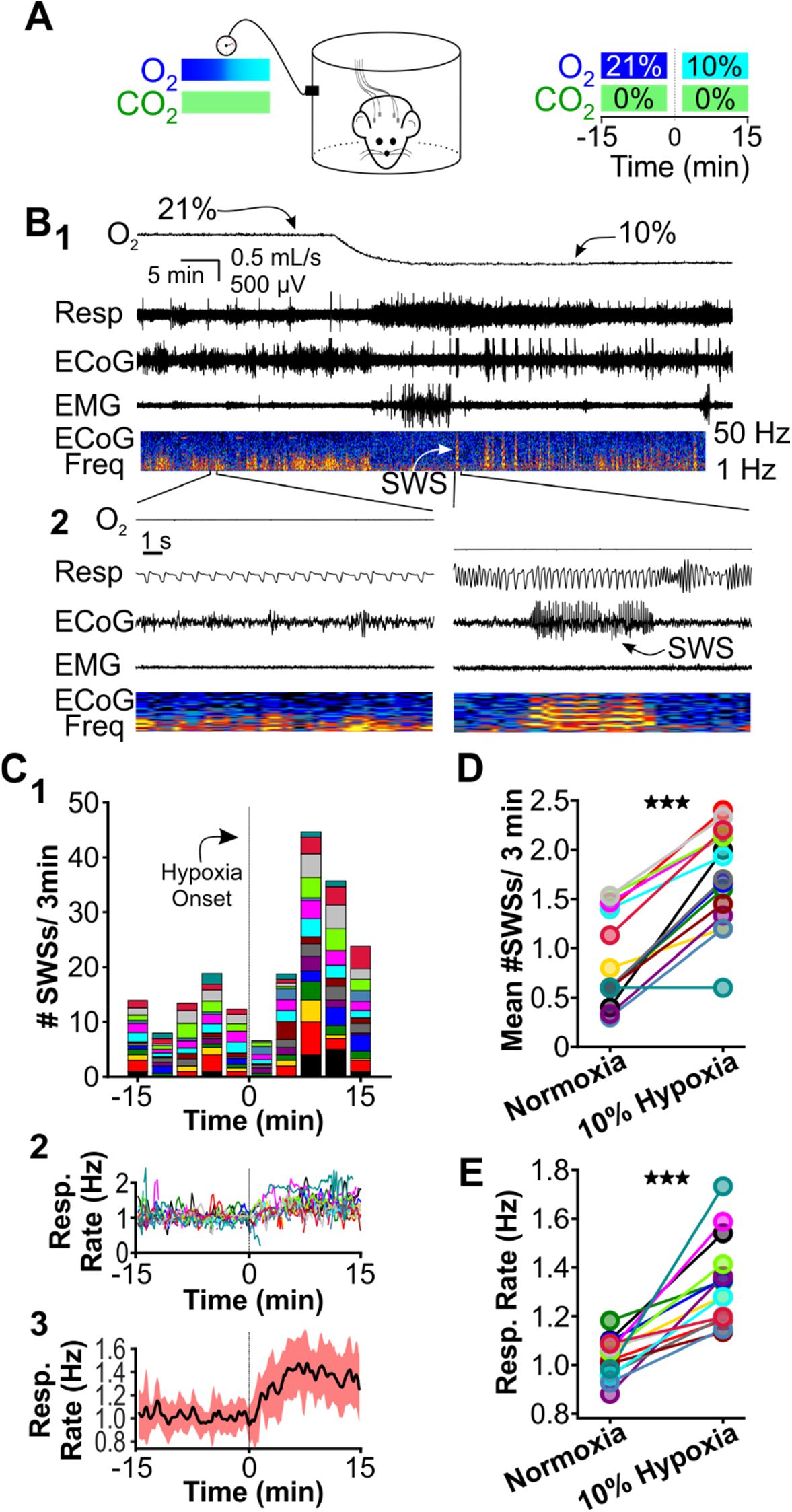
Hypoxia provokes hyperventilation-associated spike-wave seizures in WAG/Rij rats. **(A)** Experiment Paradigm. *Left:* Plethysmography chambers recorded ventilation and ECoG/EMG signals in rats exposed to normoxia (i.e., <u>21%</u> O_2_) and hypoxia (i.e., <u>10%</u> O_2_). *Right:* Example gas exchange protocol used to generate the peristimulus time histogram in panel C. Spike-wave seizure count was measured during the 15 minutes before and after gas exchange at t = 0 min. **(B)** Representative recordings during transition from normoxia to hypoxia. (1) From top to bottom: chamber O_2_, respiration, ECoG, EMG, and ECoG power spectrogram. White arrow points to spike-wave seizure. (2) *Bottom:* expanded view B1. Spectrogram reveals 5-8 Hz frequency harmonics associated with spike-wave seizures. **(C)** Spike-wave seizure (SWS) and respiration quantification. (1) Stacked histogram illustrating spike-wave seizure count for each animal before and after the onset of hypoxia; each color is a different rat. Arrow points to gas exchange at t = 0 min. (2) Corresponding respiratory rate for each animal shown in panel C1. (3) Mean respiratory rate for all animals. **(D)** Mean spike-wave seizure count per bin and **(E)** respiratory rate before and after gas exchange. See **Tables 1 & 2** for detailed statistics. ***p < 0.001.

We first compared respiration and ECoG/EMG activity in rats exposed to atmospheric conditions (i.e., normoxia: <u>21%</u> O_2_; <u>0%</u> CO_2_; <u>79%</u> N_2_) and hypoxia (<u>10%</u> O_2_; <u>0%</u> CO_2_; <u>90%</u> N_2_).

Hypoxia reliably stimulates rapid breathing, blood alkalosis and hypocapnia in rats (Basting et al., 2015; Souza et al., 2019). We cycled rats between 40-minute epochs of normoxia and 20-minute epochs of hypoxia. O_2_ levels were measured from the outflow of the plethysmography chamber for confirmation of gas exchange (Figure 1B, *top*). Hypoxia evoked a robust increase in respiratory rate (Figure 1B, *expanded*) and reliably provoked seizures. A peristimulus time histogram (PSTH) aligned to the onset of gas exchange shows spike-wave seizure counts during the 15 minutes immediately before and during hypoxia (Figure 1C1); the PSTH shows the contribution of each rat in stacked histogram format. Respiratory rates confirmed that hypoxia increased ventilation (Figure 1C2,3). To quantify the effect of hypoxia on seizures, we calculated the mean spike-wave seizure count across all bins for each rat. Relative to normoxia, spike-wave seizure count during hypoxia was nearly 2-fold higher (p = 4.5 x 10^-7^, n = 15; Fig. 1D) and respiratory rate increased by 30% (p = 1.6 x 10^-5^, n = 15; Fig. 1E).

Recent work shows that spike-wave seizures commonly occur in several rat strains, including those that are generally not considered epileptic (Taylor et al., 2017, 2019). While between 62% (Vergnes et al., 1982) and 84% (Robinson & Gilmore, 1980) of Wistar rats do not have seizures, we nonetheless tested whether hypoxia can unmask seizure-generating potential in this strain, as Wistar and WAG/Rij rats share the same genetic background (Festing, 1979). In normoxia, seizures were absent in all four Wistar rats we tested, consistent with the infrequent spike-wave seizure occurrence reported for this strain. Relative to normoxia in Wistar rats, hypoxia induced hyperventilation, hypocapnia and blood alkalization but did not provoke spike-wave seizures (Figure 2; see Table 3). Instead, hypoxia primarily triggered arousal in Wistar rats, as revealed in EEG spectrograms by the reduction in sleep-related frequencies. Therefore, we hypothesize that hypoxia-provoked spike-wave seizures are unique to seizure-prone rodent models, just as hyperventilation does not provoke absence seizures in otherwise healthy humans.

**Figure 2.**
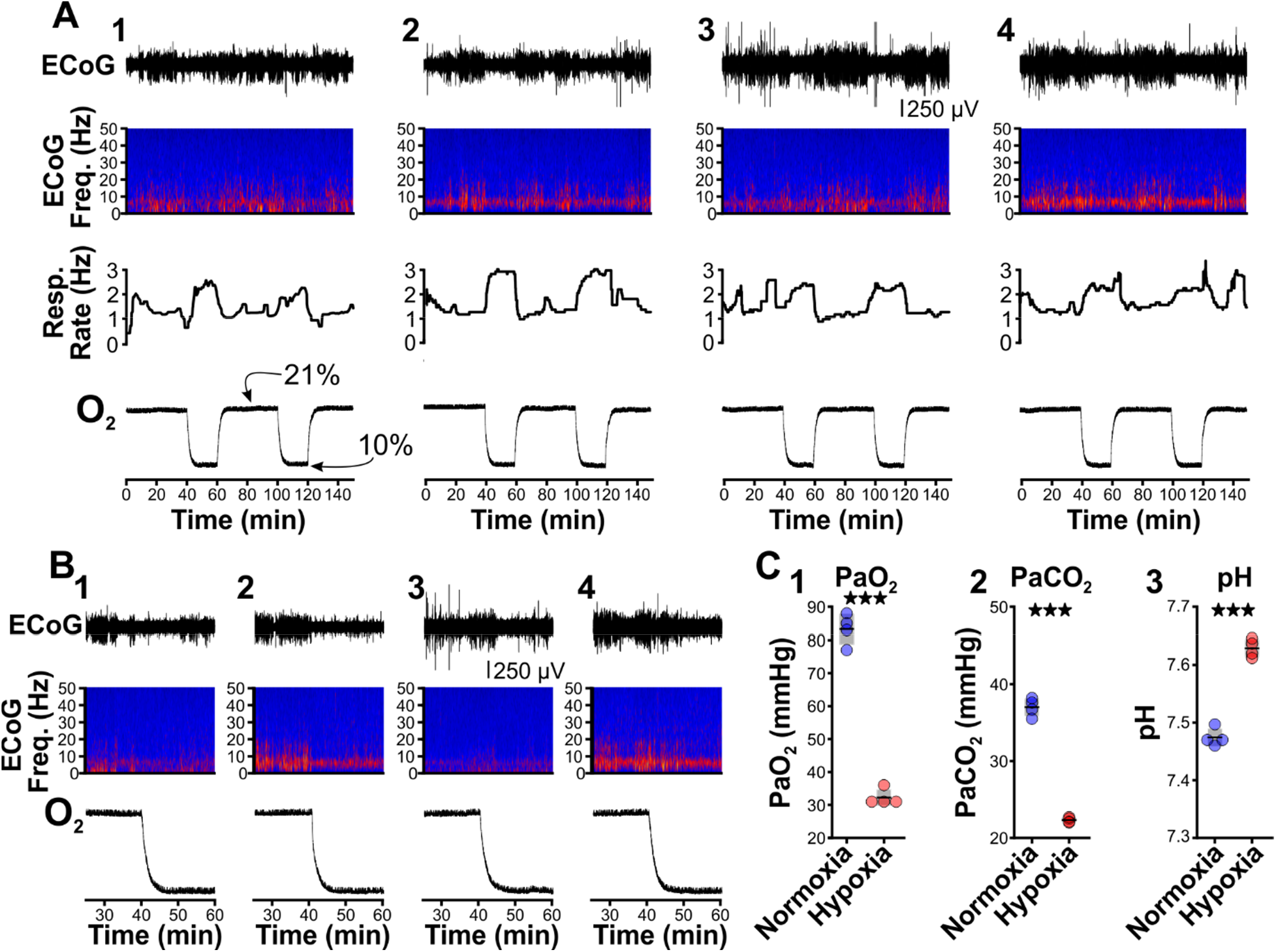
Hypoxia does not provoke hyperventilation-associated spike-wave seizures in Wistar rats. **(A)** Plethysmography chambers recorded ventilation and ECoG/EMG signals in four Wistar rats exposed to normoxia (i.e., <u>21%</u> O_2_) and hypoxia (i.e., <u>10%</u> O_2_). Panels 1-4 include responses from four Wistar rats, respectively, and show from top to bottom: ECoG, ECoG power spectrogram, respiratory rate, and chamber O_2_. During the 2.5-hour recording session, rats were challenged twice with hypoxia. No spike-wave seizures were observed during either normoxia or hypoxia. **(B)** Expanded views of the first transition from normoxia to hypoxia shown in panel A. Increased low frequency power during normoxia in some rats (e.g., panel B2) represents sleep. Hypoxia in Wistar rats generally increased arousal. **(C)** Arterial measurements in the same rats show that hypoxia challenges produced a predictable drop in arterial (1) O_2_ and (2) CO_2_, as well as (3) alkalosis. See **Table 3** for detailed statistics. ***p < 0.001.

**Table 1.**
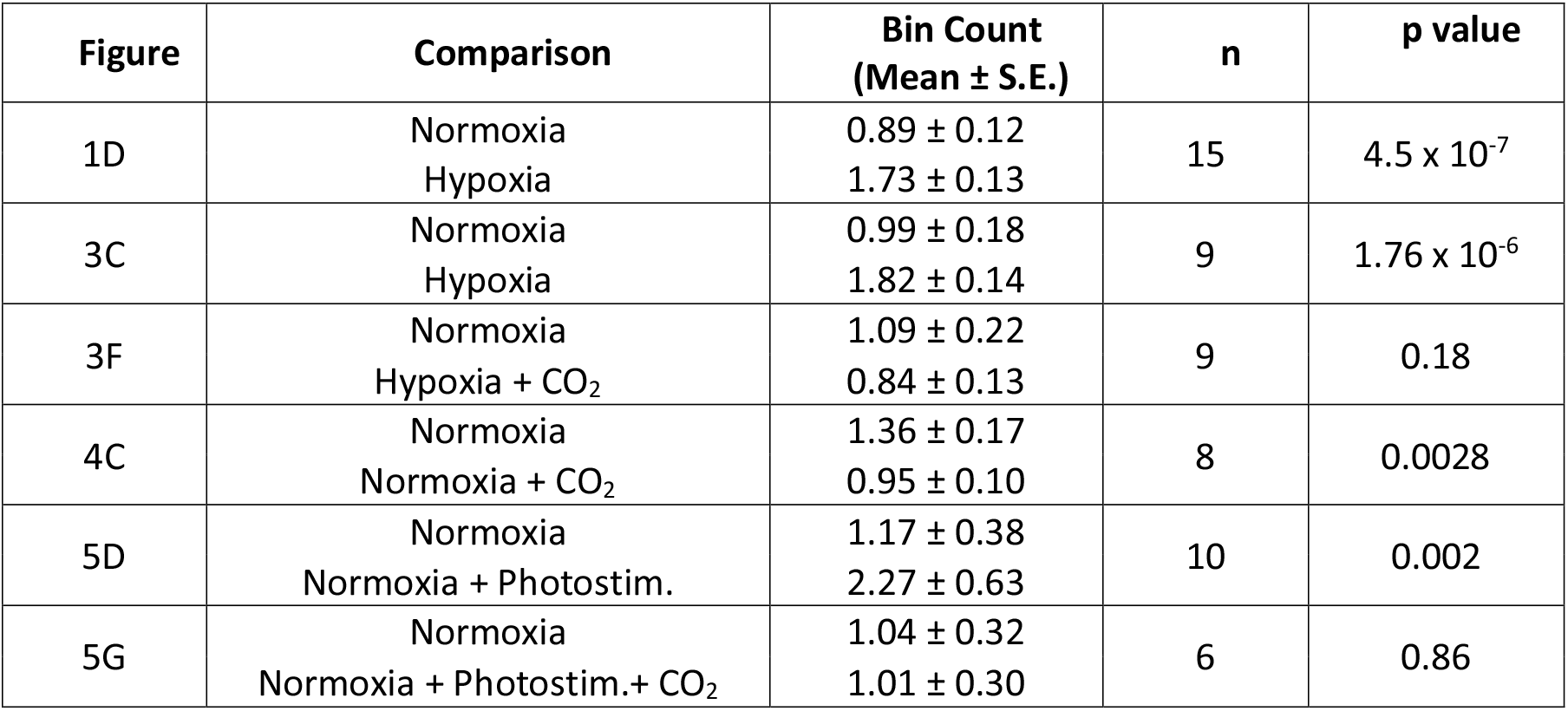
Spike-wave seizure count.

**Table 2.**
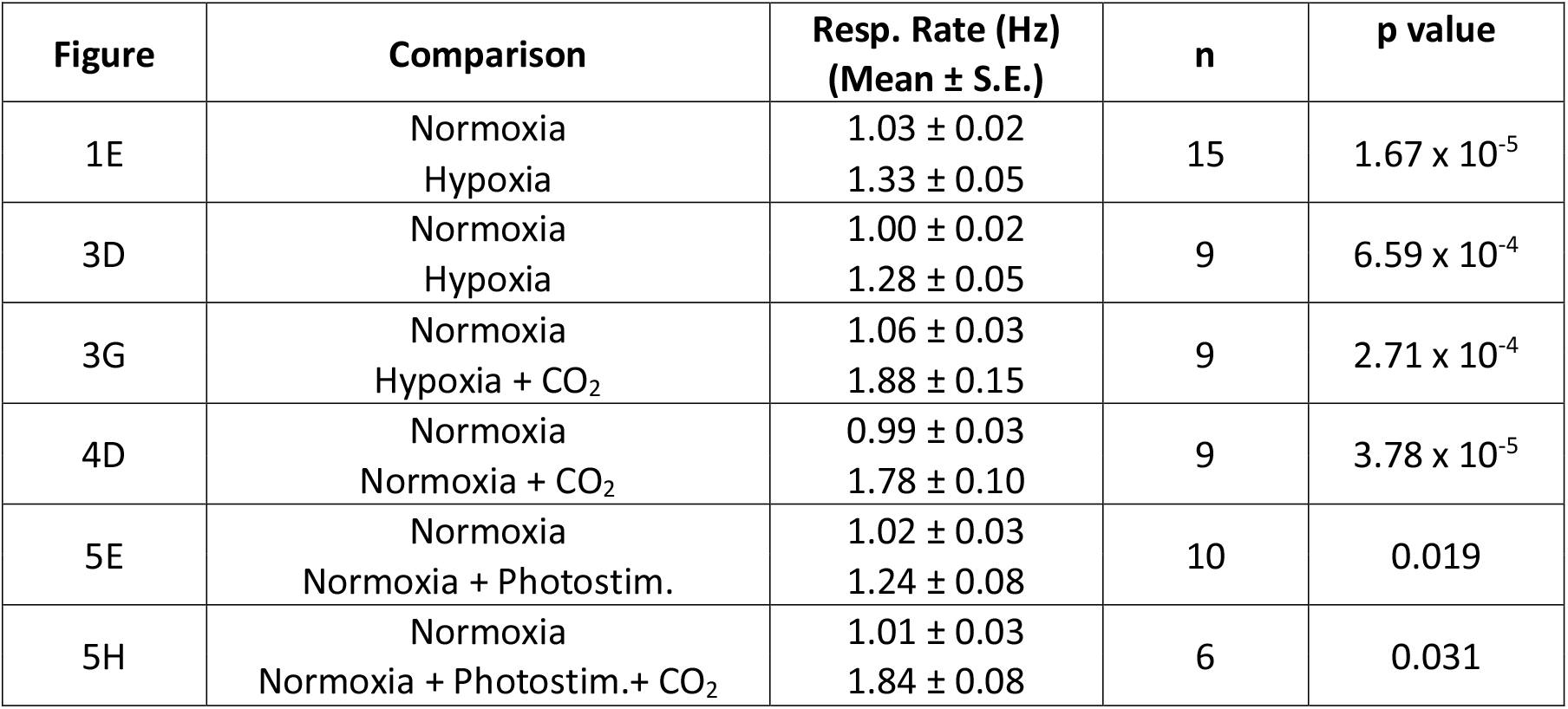
Respiratory Rate.

**Table 3.** Arterial measurements in Wistar rats.

| Figure | Parameter | Comparison | Value | n | p value |
| --- | --- | --- | --- | --- | --- |
| 2C1 | PaO<sub>2</sub> | Normoxia | 83.25 ± 2.32 | 4 | 0.0002 |
|  |  | Hypoxia | 32.25 ± 1.25 |  |  |
| 2C2 | PaCO<sub>2</sub> | Normoxia | 37.0 ± 0.59 | 4 | 6.6 x 10<sup>-5</sup> |
|  |  | Hypoxia | 22.33 ± 0.16 |  |  |
| 2C3 | pH | Normoxia | 7.47 ± 0.01 | 4 | 4.5 x 10<sup>-5</sup> |
|  |  | Hypoxia | 7.63 ± 0.01 |  |  |

### CO_2_ suppresses spike-wave seizures

Hyperventilation promotes hypocapnia, a state of low PaCO_2_. As dissolved CO_2_ is acidic, hyperventilation-triggered hypocapnia is also associated with respiratory alkalosis. To test the hypothesis that hypocapnia specifically provokes seizures, we next determined whether supplemental CO_2_ (5%) blunts the spike-wave seizure-provoking effects of hypoxia. We performed ECoG/plethysmography experiments as before but alternated between two test trials: hypoxia and hypoxia/hypercapnia (<u>10%</u> O_2_, <u>5%</u> CO_2_; <u>85%</u> N_2_). Test trials were interleaved with 40-minute periods of normoxia to allow blood gases to return to baseline levels (Figure 3A). As before, hypoxia increased spike-wave seizure count by nearly 2-fold (p = 1.76 x 10^-6^, n = 9; Figure 3B1, C) and increased respiratory rate by 27% (p = 6.59 x 10^-4^, n = 9; Figure 3B3, D). In the same rats, supplementing hypoxia with 5% CO_2_ suppressed the spike-wave seizure response insofar that hypoxia/hypercapnia did not change spike-wave seizure count relative to normoxia (p = 0.18, n = 9; Figures 3E1 and 3F) despite a predictable and robust elevation in respiratory rate (p = 2.71 x 10^-4^, n = 9; Figures 3E2, 3 and 3G).

**Figure 3.**
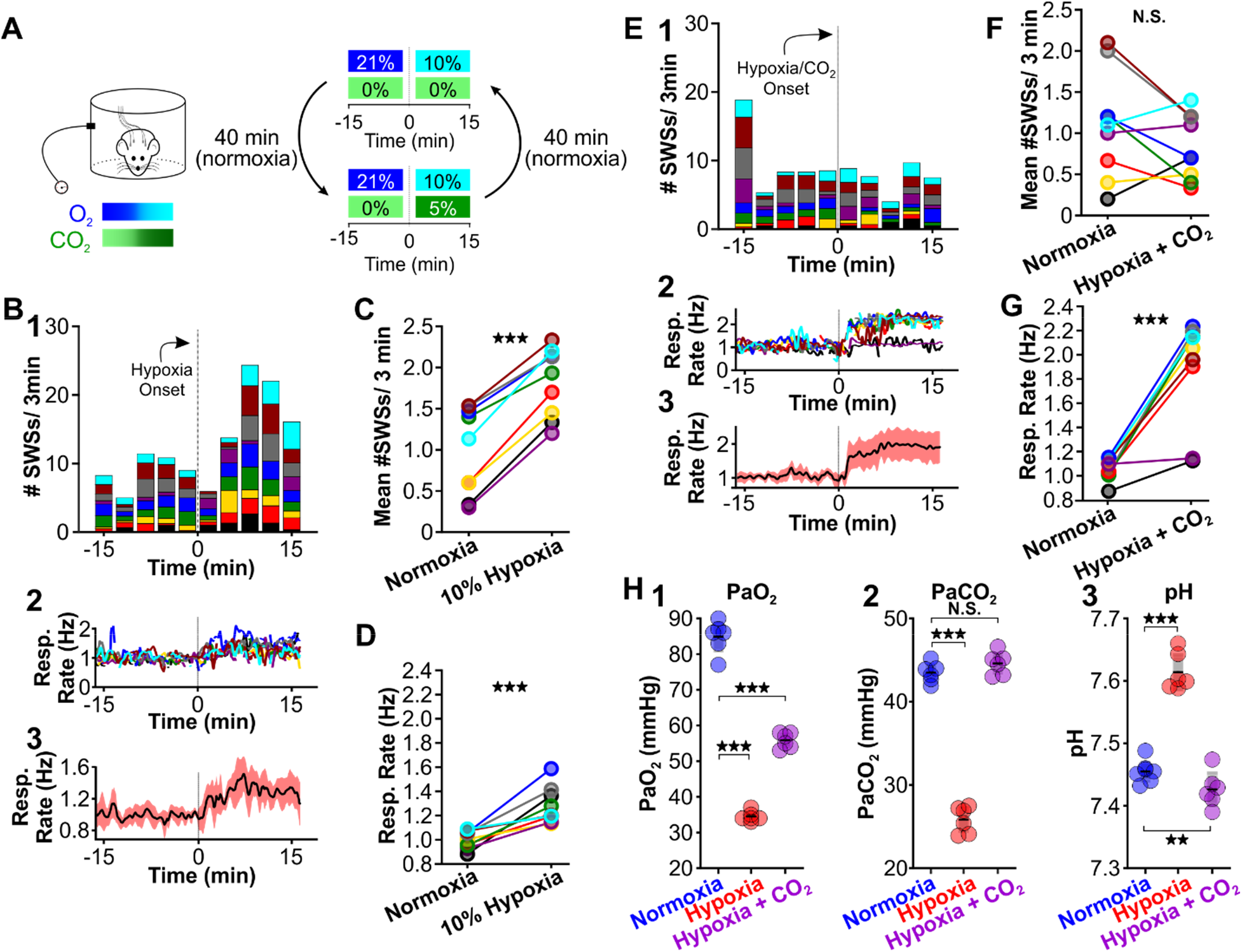
Supplemental CO_2_ suppresses hypoxia-provoked spike-wave seizures. **(A)** Experimental approach. Plethysmography chambers recorded ventilation and ECoG/EMG signals in WAG/Rij rats exposed to normoxia (i.e., <u>21%</u> O_2_) and then alternately challenged with hypoxia (i.e., <u>10%</u> O_2_) or hypoxia + CO_2_, (i.e., <u>10%</u> O_2_, <u>5%</u> CO_2_). **(B-D)** Hypoxia challenge. **(B)** Spike-wave seizure (SWS) and respiration quantification. (1) Stacked histogram illustrating spike-wave seizure count for each animal before and after the onset of hypoxia. (2) Corresponding respiratory rate for each animal shown in panel B1. (3) Mean respiratory rate for all animals. **(C)** Mean spike-wave seizure count per bin and **(D)** respiratory rate before and after hypoxia exchange. **(E-G)** Hypoxia + CO_2_ challenge. **(E)** SWS and respiration quantification. (1) Stacked histogram illustrating spike-wave seizure count for each animal before and after the onset of hypoxia + CO_2_. (2) Corresponding respiratory rate for each animal shown in panel E1. (3) Mean respiratory rate for all animals. **(F)** Mean spike-wave seizure count per bin and **(G)** respiratory rate before after hypoxia + CO_2_ exchange. **(H)** Arterial measurements in the same rats show that hypoxia produced a predictable drop in arterial (1) O_2_ and (2) CO_2_, as well as (3) respiratory alkalosis (as in Wistar rats). Supplementing the chamber with 5% CO_2_ normalizes arterial CO_2_ and pH. Elevated arterial O_2_ during hypoxia + CO_2_ relative to hypoxia reflects a powerful inhalation response during the former condition (c.f., panels D and G). See **Tables 1, 2 and 4** for detailed statistics. **p<0.01, ***p < 0.001.

In a separate cohort of rats, we collected arterial blood samples to measure blood PaCO_2_, PaO_2_ and pH during normoxia, hypoxia and hypoxia/hypercapnia (see Table 4). We observed a considerable change in PaO_2_ [F (1.056, 5.281) = 406.4, p = 3.0 x 10^-6^], PaCO_2_ [F (1.641, 8.203) = 338.9, p = 1.9 x 10^-8^] and pH [F (1.938, 9.688) = 606, p = 7.2 x 10^-11^] values among the three conditions. Hypoxia decreased PaCO_2_ (p = 2.1 x 10^-6^; n = 6; Figure 3H2) and concomitantly alkalized the blood (p = 7.0 x 10^-6^, n = 6; Figure 3H3). We also observed a decrease in PaO_2_ (p = 6.0 x 10^-6^; n = 6; Figure 3H1). Supplemental CO_2_ returned blood pH (p = 0.008, n = 6; Figure 3H3) and PaCO2 (p = 0.42, n = 6; Figure 3H2) to normoxia levels. However, heightened respiratory rate in supplemental CO_2_ raised PaO_2_ (p = 00013, n = 6; Figure 3H1). Collectively, these data support the hypothesis that blood pH powerfully regulates spike-wave seizure activity.

**Table 4.** Arterial measurements in WAG/Rij rats.

| Figure | Parameter | Comparison | Value | n | p value |
| --- | --- | --- | --- | --- | --- |
| 3H1 | PaO<sub>2</sub> | Normoxia | 84.93 ± 1.82 | 6 | 6.0 x 10<sup>-6</sup> |
|  |  | Hypoxia | 34.50 ± 0.56 |  |  |
|  |  | Normoxia | 84.93 ± 0.02 | 6 | 0.000134 |
|  |  | Hypoxia +CO<sub>2</sub> | 55.83 ± 0.87 |  |  |
| 3H2 | PaCO<sub>2</sub> | Normoxia | 43.48 ± 0.47 | 6 | 2.1 x 10<sup>-6</sup> |
|  |  | Hypoxia | 25.83 ± 0.65 |  |  |
|  |  | Normoxia | 43.48 ± 0.47 | 6 | 0.42 |
|  |  | Hypoxia +CO<sub>2</sub> | 44.60 ± 0.55 |  |  |
| 3H3 | pH | Normoxia | 7.45 ± 0.01 | 6 | 7.0 x 10<sup>-6</sup> |
|  |  | Hypoxia | 7.61 ± 0.01 |  |  |
|  |  | Normoxia | 7.45 ± 0.01 | 6 | 0.008 |
|  |  | Hypoxia +CO<sub>2</sub> | 7.43 ± 0.01 |  |  |
| 4E1 | PaO<sub>2</sub> | Normoxia | 84.93 ± 1.82 | 6 | 0.00019 |
|  |  | 5% CO<sub>2</sub> | 34.50 ± 0.56 |  |  |
| 4E2 | PaCO<sub>2</sub> | Normoxia | 43.48 ± 0.47 | 6 | 0.022 |
|  |  | 5% CO<sub>2</sub> | 25.83 ± 0.65 |  |  |
| 4E3 | pH | Normoxia | 7.45 ± 0.01 | 6 | 0.00063 |
|  |  | 5% CO<sub>2</sub> | 7.42 ± 0.01 |  |  |

**Table 5.** cFos-positive cells in WAG/Rij rats.

| Figure | Threshold | Comparison | Counts<br>(Mean ± S.E.) | n | p value |
| --- | --- | --- | --- | --- | --- |
| 6C | 3 | Normoxia<br>Hypoxia | 282 ± 148.2<br>1370 ± 137 | 4 | 1.5 × 10<sup>-7</sup> |
| | | Normoxia<br>Hypoxia + CO2 | 282 ± 148.2<br>385.5 ± 78.7 | 4 | 0.55 |
| | | Hypoxia<br>Hypoxia + CO2 | 1370 ± 137<br>385.5 ± 78.7 | 4 | 4.3 × 10<sup>-7</sup> |
| | 5 | Normoxia<br>Hypoxia | 112.3 ± 57.1<br>595.3 ± 85.0 | 4 | 0.0005 |
| | | Normoxia<br>Hypoxia + CO2 | 112.3 ± 57.1<br>348 ± 68.9 | 4 | 0.045 |
| | | Hypoxia<br>Hypoxia + CO2 | 595.3 ± 85.0<br>348 ± 68.9 | 4 | 0.061 |
| | 7 | Normoxia<br>Hypoxia | 57.3 ± 29.2<br>349 ± 75.0 | 4 | 0.021 |
| | | Normoxia<br>Hypoxia + CO2 | 57.3 ± 29.2<br>319.5 ± 63.1 | 4 | 0.036 |
| | | Hypoxia<br>Hypoxia + CO2 | 349 ± 75.0<br>319.5 ± 63.1 | 4 | 0.95 |

Next, we tested whether supplementing normoxia with 5% CO_2_ is sufficient to reduce spike-wave seizure counts. Respiration during high CO_2_ causes hypercapnia, a condition that increases blood PaCO_2_ and acidifies the blood (Eldridge et al., 1984). As with hypoxia, hypercapnia also triggers hyperventilation (Guyenet et al., 2019). We performed ECoG/plethysmography experiments in rats that cycled through trials of normoxia and hypercapnia (<u>21%</u> O_2_; <u>5%</u> CO_2_; <u>74%</u> N_2_) and compared the mean number of seizures observed during the two conditions. Relative to normoxia, the number of spike-wave seizures was lower during 5% CO_2_ (p = 0.0028, n = 8; Figure 4B1 and 4C); hypercapnia also induced a powerful respiratory response (p = 3.78 x 10^-5^, n = 8; Figure 4B2,3 and 4D). Blood gas measurements revealed that 5% hypercapnia increased PaCO_2_ (p = 0.022, n = 6; Figure 4E2) and slightly acidified blood pH (p = 0.00063, n = 6; Figure 4E3). These results provide further support for the hypothesis that the neural circuits that produce spike-wave seizures are CO_2_-sensitive, and thus pH-sensitive. Moreover, the results demonstrate that neither the mechanics of elevated ventilation, nor increased arousal, is sufficient to provoke spike-wave seizures.

**Figure 4.**
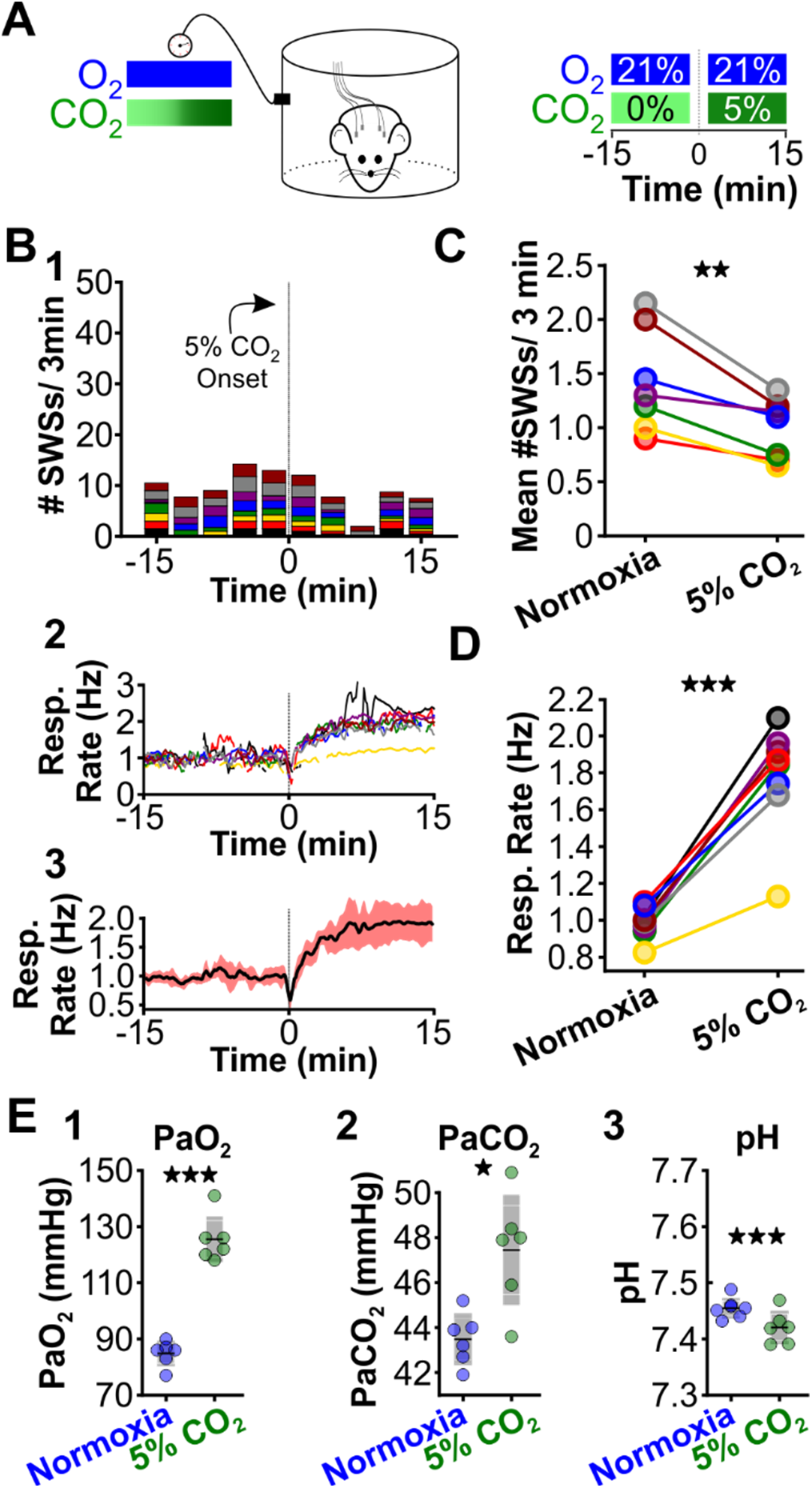
Supplemental CO_2_ suppresses spontaneous spike-wave seizures. **(A)** Experimental approach. Plethysmography chambers recorded ventilation and ECoG/EMG signals in WAG/Rij rats exposed to normoxia (i.e., <u>21%</u> O_2_) and hypercapnia (i.e., <u>21%</u> O_2_, <u>5%</u> CO_2_). **(B)** Spike-wave seizure (SWS) and respiratory quantification. (1) Stacked histogram illustrating spike-wave seizure count for each animal before and after the onset of hypercapnia. (2) Corresponding respiratory rate for each animal shown in panel B1. (3) Mean respiratory rate for all animals. **(C)** Mean spike-wave seizure count per bin and **(D)** respiratory rate before and after hypercapnia exchange. **(E)** Arterial measurements in the same rats show that hypercapnia produced a predictable increase in arterial (1) O_2_ and (2) CO_2_, as well as (3) respiratory acidosis. Increase arterial O_2_ reflects robust ventilatory response during hypercapnia. See **Tables 1, 2 and 4** for detailed statistics. *p < 0.05, **p < 0.01, ***p < 0.001.

### Optogenetic stimulation of the retrotrapezoid nucleus provokes spike-wave seizures

In addition to inducing hyperventilation and hypocapnia, hypoxia also lowers PaO_2_ (see Figure 3H1), an effect that stimulates the carotid body, the principal peripheral chemoreceptor that initiates hyperventilation during hypoxic conditions (Lindsey et al., 2018; Lopez-Barneo et al., 2016; Semenza & Prabhakar, 2018). Carotid body activity recruits neurons of the nucleus tractus solitarius (NTS) that then excite neurons of the central respiratory pattern generator to drive a respiratory response (Guyenet, 2014; Lopez-Barneo et al., 2016). To evaluate the capacity of hyperventilation to provoke seizures in the absence of hypoxia (and, therefore, in the absence of carotid body activation), we utilized an alternative approach to induce hyperventilation. Under physiological conditions, chemosensitive neurons of the retrotrapezoid nucleus (RTN), a brainstem respiratory center, are activated during an increase in PaCO_2_ and a consequent drop in arterial pH (Guyenet et al., 2016, 2019; Guyenet & Bayliss, 2015) that then stimulate respiration. Optogenetic activation of RTN neurons in normoxia is sufficient to evoke a powerful hyperventilatory response that alkalizes the blood (Abbott et al., 2011; Souza et al., 2020). Importantly, PaO_2_ remains stable (or is slightly elevated) during optogenetically-induced respiration. Therefore, hyperventilation evoked by optogenetic RTN activation during normoxia both (1) promotes respiratory alkalosis without hypoxia and (2) is a more clinically relevant approximation of voluntary hyperventilation than hypoxia-induced hyperventilation.

We selectively transduced RTN neurons of WAG/Rij rats with a lentiviral approach using the PRSX8 promoter to drive channelrhodopsin expression (Abbott et al., 2009; Hwang et al., 2001; Lonergan et al., 2005). Once channelrhodopsin was expressed, we challenged rats with two test trials: RTN photostimulation during normoxia and RTN photostimulation during hypercapnia (Figure 5A); in a subset of animals, we cycled rats between the two conditions. In both trials, the laser was pulsed at 20 Hz (10msec pulse) once every four seconds for two seconds. Laser stimulation during normoxia provoked spike-wave seizures (p = 0.002; n = 10; Figures 5B, 5C1 and 5D) and also increased ventilation (p = 0.019; n = 10; Figures 5C2,3 and 5E). In contrast, laser stimulation during hypercapnia in the same animals did not alter spike-wave seizure count (p = 0.86; n = 6; Figures 5F1 and 5G), despite the induction of a strong hyperventilatory response (p = 0.031; n = 6; Figures 5F2,3 and 5H). In sum, these results support the hypothesis that respiratory alkalosis is necessary to provoke seizures during hyperventilation and excludes carotid body activation as a contributing factor.

**Figure 5.**
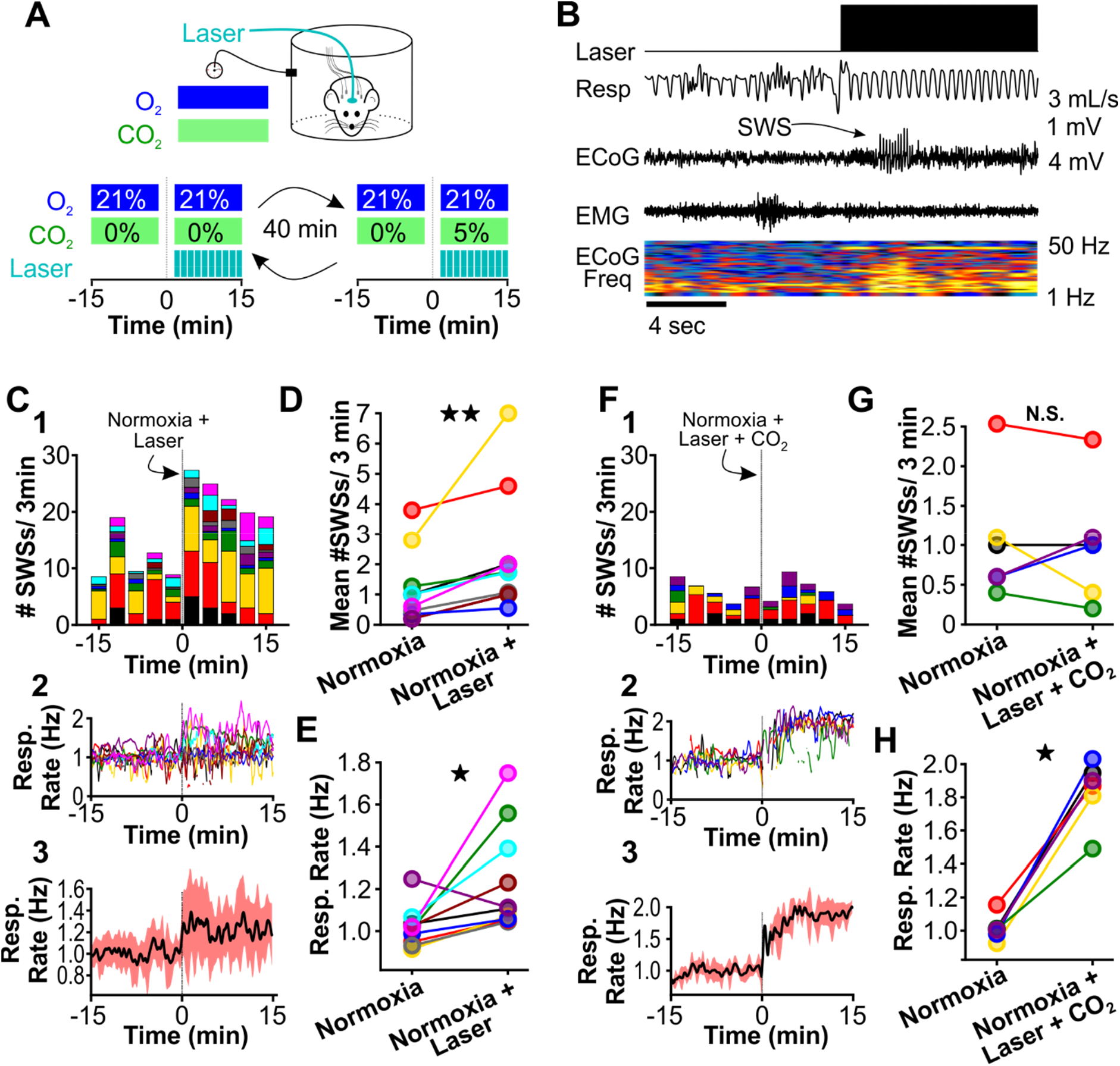
Normoxic hyperventilation provokes CO_2_-sensitive spike-wave seizures. **(A)** Experimental approach. Plethysmography chambers recorded ventilation and ECoG/EMG signals in WAG/Rij rats exposed to normoxia (i.e., <u>21%</u> O_2_) and normoxia + CO_2_, (i.e., <u>10%</u> O_2_, <u>5%</u> CO_2_). Channelrhodopsin-mediated photostimulation of the retrotrapezoid nucleus (RTN) was used to increase ventilation. **(B)** Example of ventilatory response and spike-wave seizure during normoxic RTN photostimulation **(C-E)** RTN photostimulation during normoxia. **(C)** Spike-wave seizure (SWS) and respiration quantification. (1) Stacked histogram illustrating spike-wave seizure count for each animal before and after normoxia photostimulation onset. (2) Corresponding respiratory rate for each animal shown in panel C1. (3) Mean respiratory rate for all animals. **(D)** Mean spike-wave seizure count per bin and **(E)** respiration rate before and after normoxia photostimulation onset. **(F-H)** RTN photostimulation during hypercapnia (i.e., <u>21%</u> O_2_, <u>5%</u> CO_2_). **(F)** Spike-wave seizure and respiratory quantification. (1) Stacked histogram illustrating spike-wave seizure count for each animal before and after hypercapnic photostimulation onset. (2) Corresponding respiratory rate for each animal shown in panel F1. (3) Mean respiratory rate for all animals. **(G)** Mean spike-wave seizure count per bin and **(H)** respiratory rate before and after hypercapnic photostimulation onset. See **Tables 1, 2 and 4** for detailed statistics. *p < 0.05, **p < 0.01, not significant (n.s.).

### Hypoxia-induced hyperventilation activates neurons of the intralaminar thalamus

Thus far, our results demonstrated that respiratory alkalosis (i.e., hyperventilation that promotes a net decrease in PaCO_2_) provokes spike-wave seizures in the WAG/Rij rat. Next, we sought to identify brain structures activated during respiratory alkalosis that may contribute to spike-wave seizure provocation. We used the neuronal activity marker cFos to identify such structures in WAG/Rij rats. To isolate activation specifically associated with respiratory alkalosis, we first administered ethosuximide (200mg/kg, i.p.) to suppress spike-wave seizures; respiration and ECoG/EMG signals confirmed ventilatory responses and spike-wave seizure suppression. Ethosuximide-injected rats were exposed to either hypoxia, normoxia or hypoxia/hypercapnia for 30 minutes and then transcardially perfused 90 minutes later. Brains were harvested and evaluated for cFos immunoreactivity. Surprisingly, in rats exposed to hypoxia we observed heightened immunoreactivity in the intralaminar nuclei, a group of higher-order thalamic nuclei that, unlike first-order thalamic nuclei, do not receive peripheral sensory information (Saalmann, 2014) (Figure 6A,B). Indeed, cFos immunoreactivity was largely absent from first-order thalamic nuclei and cortex, and was blunted in rats treated with normoxia and hypoxia/hypercapnia (Figure 6B). Importantly, the latter condition elevates respiration but normalizes arterial pH (see Figure 3G and 3H). Immunoreactivity quantification revealed that the number of cFos-positive cells within the intralaminar thalamic nuclei was highest following hypoxia [ANOVA: F (2, 6) = 31.59, p = 0.00019, Figure 6C].

**Figure 6.**
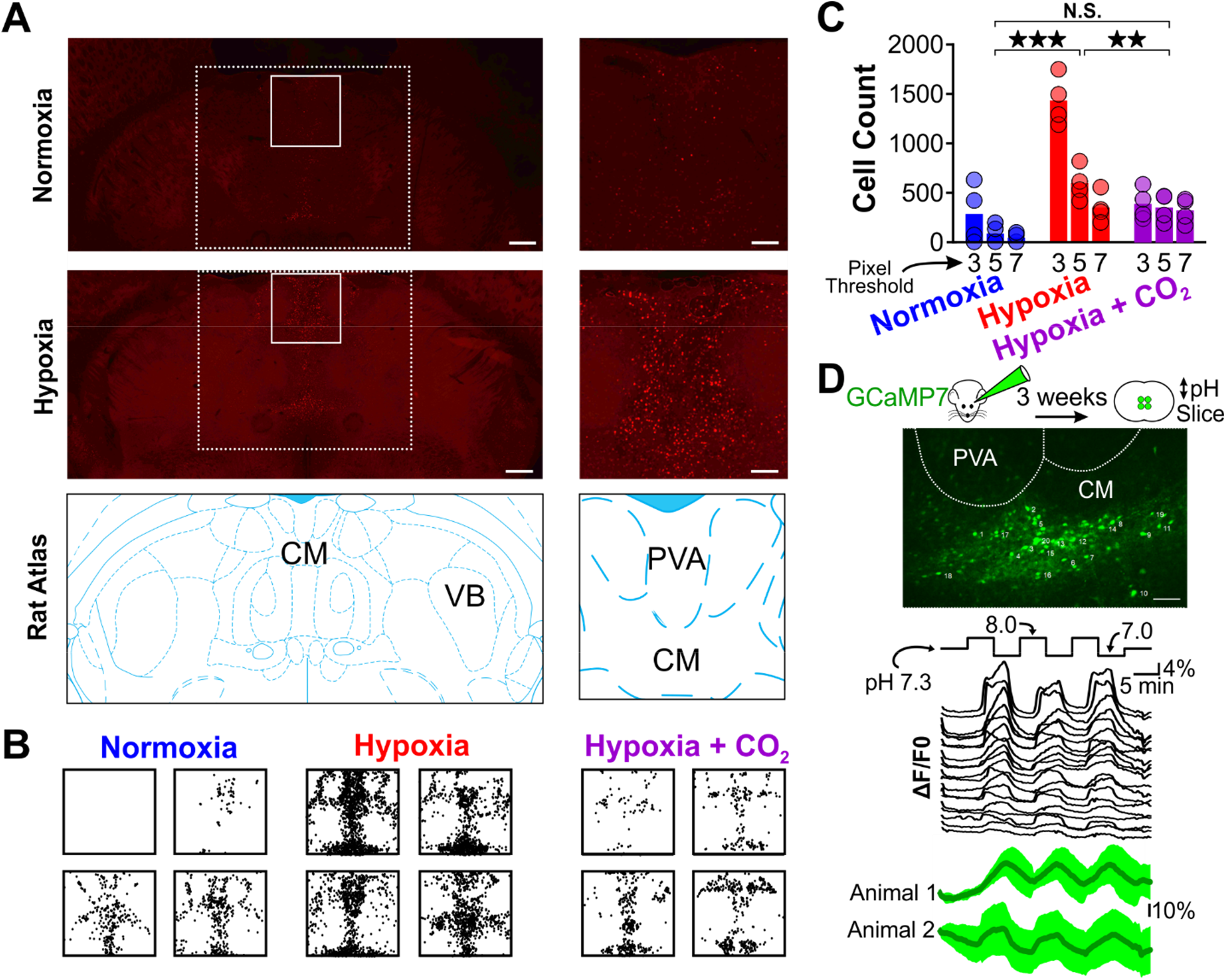
Hypoxia-induced hyperventilation activates intralaminar thalamic neurons. **(A)** cFos immunohistochemistry in horizontal sections of the WAG/Rij rat. Dashed lines highlight the medial region of the thalamus containing the intralaminar nuclei. Solid lines demarcate regions containing elevated cFos expression and are expanded on right. Top images are collected from a rat exposed to 30 minutes of normoxia. Middle images are collected from a rat exposed to 30 minutes of hypoxia. Bottom images are taken from Paxinos and Watson (Paxinos & Watson, 2007) and show the structural landmarks in the top and middle images. The central median nucleus (CM, intralaminar thalamus) and ventrobasal complex (VB, first-order thalamus) are labeled. **(B)** cFos density plots show immunoreactivity in each of four rats exposed to either normoxia, hypoxia or hypoxia + CO_2_. Each black dot represents a cFos-positive cell, as identified with ImageJ (see Methods). Plots are aligned to expanded views in panel A. **(C)** Quantification of cFos labeled cells at different ImageJ thresholding values. **(D)** GCaMP7 was stereotaxically delivered to the intralaminar nuclei. Later, fluorescence changes were measured during extracellular alkaline challenges in acute slices containing the intralaminar nuclei. Individual ROIs show fluorescence changes during alkalosis (black traces). Mean responses from two animals are shown in green. **p < 0.01, ***p < 0.001. See **Table 5** for detailed statistics. Scale bars are 500 pm (*left*) and 100 pm (*right*).

As heightened cFos immunoreactivity was observed primarily following hypoxia that results in pronounced respiratory alkalosis, we next tested the hypothesis that neurons of the intralaminar nuclei are pH-sensitive. We stereotaxically delivered the pan-neuronal expressing GCaMP7s (pGP-AAV-syn-jGCaMP7s-WPRE) to the intralaminar nuclei and harvested acute brain sections three weeks later (Figure 6D). Recording fluorescence changes in brain sections revealed that extracellular alkalosis quickly and reversibly activated neurons of the intralaminar nuclei (Figure 6D). Collectively, these results support the hypothesis that respiratory alkalosis activates pH-sensitive neurons of the intralaminar thalamic nuclei in the WAG/Rij rat.

## Discussion

Hyperventilation-provoked seizures associated with absence epilepsy were first formally described in 1928 by William Lennox (Lennox, 1928) and despite the clinical ubiquity of utilizing hyperventilation to diagnose the common form of childhood epilepsy, no animal studies have attempted to resolve the physiological events that enable hyperventilation to reliably provoke spike-wave seizures. To resolve events and relevant brain structures recruited during this phenomenon, we first utilized the WAG/Rij rat to establish a rodent model that mimics hyperventilation-provoked spike-wave seizures in humans. With this model, we show that hyperventilation only provokes spike-wave seizures in seizure-prone, not generally seizure-free, rats. We then show that supplemental CO_2_, by mitigating respiratory alkalosis, suppresses spike-wave seizures triggered by hyperventilation during either hypoxia or direct activation of brainstem respiratory centers. Moreover, supplemental CO_2_, by producing respiratory acidosis, suppresses spontaneous spike-wave seizures (i.e., those occurring during normoxia) despite a compensatory increase in respiratory rate. These data demonstrate that spike-wave seizures are yoked to arterial CO_2_/pH. Finally, we demonstrate that respiratory alkalosis activates neurons of the intralaminar thalamic nuclei, also in a CO_2_-dependent manner; activation of these neurons is also pH-sensitive. With these observations, we propose a working model wherein respiratory alkalosis activates pH-sensitive neurons of the intralaminar nuclei that in turn engage seizure-generating neural circuits to produce spike-wave seizures (Figure 7).

**Figure 7.**
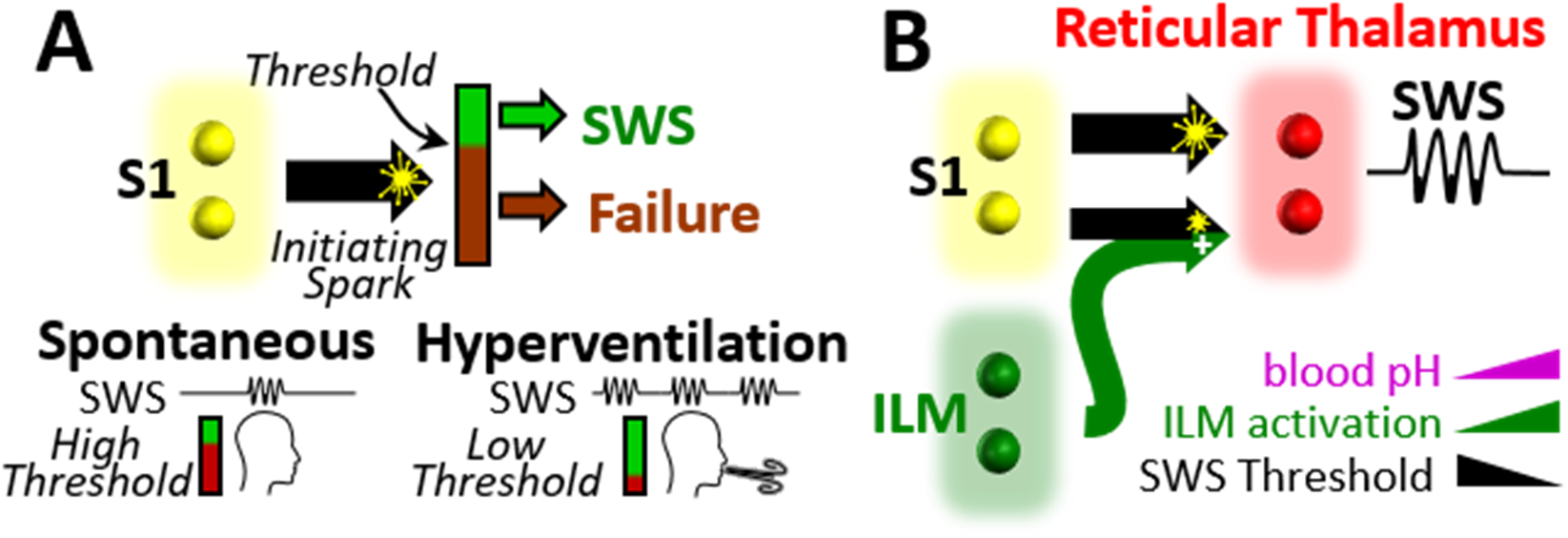
Working model. **A.** Spike-wave seizures only occur if initiating activity from S1 somatosensory cortex successfully overcomes a threshold, consistent with the cortical focus theory (H. K. M. Meeren et al., 2002). Hyperventilation-associated alkalosis reduces spike-wave seizure (SWS) threshold. **B.** S1 initiating activity is proposed to overcome a seizure node formed by circuits in reticular thalamus to generate an spike-wave seizure (Paz & Huguenard, 2015). We propose that hyperventilation-evoked respiratory alkalosis activates the intralaminar nuclei (ILM) to reduce the threshold for S1 activity required to evoke a spike-wave seizure.

### Cortical EEG Patterns Evoked by Hyperventilation

Hyperventilation produces stereotypical EEG patterns in both healthy children and children with absence epilepsy (Barker et al., 2012). In healthy children, hyperventilation can evoke an EEG pattern known as *Hyperventilation-Induced, High-Amplitude Rhythmic Slowing* (HIHARS) that is often associated with altered awareness (Barker et al., 2012; Lum et al., 2002). Electrographically, HIHARS is distinct from spike-wave seizures insofar the EEG lacks epilepsy-associated spikes and resembles slow-wave sleep. Nonetheless, similarities between HIHARS and absence seizures exist. Both events are associated with children of the same age (Mattozzi et al., 2021). Behaviorally, eye opening/staring and fluttering, as well as oral automatisms, are observed during both events, albeit with different frequencies (Lum et al., 2002). Finally, the mean latencies from the onset of hyperventilation to the onset of electrographic HIHARS in healthy children, or spike-wave seizures in absence patients, are also similar (Lum et al., 2002; Mattozzi et al., 2021).

Recent work suggests that spike-wave seizures may limit or preclude the generation of HIHARs in children with absence epilepsy, thereby supporting the hypothesis that HIHARS and spike-wave seizures borrow from similar neural circuit mechanisms (Mattozzi et al., 2021). In this model, hyperventilation engages brain structures that initiate and/or support widespread, synchronous cortical activity. However, the trajectory of this engagement ultimately bifurcates such that either HIHARS or a spike-wave seizure is produced, but not both. When viewed alongside work performed in the 1960s by Ira Sherwin (Sherwin, 1965, 1967), our results support this model. Sherwin demonstrated that hyperventilation evokes HIHARS in cats (Sherwin, 1965), and that the stereotyped EEG pattern requires an intact central lateral nucleus of the thalamus (Sherwin, 1967). Together with the central medial (CM) and paracentral thalamic nuclei, the central lateral nucleus belongs to the anterior group of the intralaminar nuclei (Saalmann, 2014), the location of cFos immunoreactivity associated with respiratory alkalosis and pH-sensitivity (Figure 6). Indeed, at the time Sherwin postulated that the intralaminar nuclei of the thalamus are both chemoreceptive and capable of engaging widespread cortical activity (Sherwin, 1967). We now postulate that these nuclei are also instrumental for provoking spike-wave seizures during hyperventilation. If true, then resolving how and where the mechanisms that produce HIHARS diverge from those that produce spike-wave seizures becomes a central goal.

### Thalamocortical circuit involvement in spike-wave seizures

Decades of work have culminated in a canonical model wherein interconnected circuits between the cortex and thalamus support the initiation and maintenance of generalized spike-wave seizures (Avoli, 2012; Beenhakker & Huguenard, 2009; Huguenard & McCormick, 2007; McCafferty et al., 2018; McCormick & Contreras, 2001; Meeren et al., 2002). By recording from multiple sites in the WAG/Rij rat, Meeren et al. (Meeren et al., 2002) concluded that the peri-oral region of somatosensory cortex provides the bout of hypersynchronous activity that initiates a spike-wave seizure. This activity then rapidly recruits additional somatosensory cortices and the lateral dorsal thalamus, a higher-order thalamic nucleus involved in spatial learning and memory (Bezdudnaya & Keller, 2008). Finally, first-order thalamic nuclei that encode somatosensory information (i.e., the ventrobasal complex) are recruited. This stereotyped succession of events occurs within the first 500 milliseconds of the spike-wave seizure, after which the temporal relationships among cortical and thalamic structures are more unpredictable (Meeren et al., 2002). Additional studies support the hypothesis that cortical hyperactivity initiates spike-wave seizures (Pinault, 2003; Pinault et al., 1998) and have motivated what is generally referred to as the *cortical focus theory* for spike-wave seizure initiation (Meeren et al., 2005).

While resolving how seizures initiate and propagate through brain structures is of critical importance, this understanding does not necessarily address the mechanisms that drive the highly rhythmic and hypersynchronous activity associated with ongoing spike-wave seizures. Extensive work on acute brain slice preparations clearly demonstrates that circuits between first-order thalamic nuclei and the reticular thalamic nucleus are sufficient to sustain rhythmic network activities, including those comparable to absence seizures (Bal et al., 1995; Bal & McCormick, 1993; McCormick & Contreras, 2001; von Krosigk et al., 1993). In this model, feedforward inhibition provided by reticular neurons evokes robust, hypersynchronous post-inhibitory rebound bursts among thalamocortical neurons that then relay activity back to reticular thalamus and to cortex. Reticular neuron-mediated feedforward inhibition of thalamocortical neurons, coupled with reciprocal excitation from thalamocortical neurons to reticular neurons, forms the basis of a rhythmogenic circuit that is proposed to maintain spike-wave seizures. While this model very likely accounts for rhythmicity in the acute brain slice preparation, it is becoming less clear how first-order thalamocortical neurons actively contribute to the maintenance of spike-wave seizures recorded *in vivo* (Huguenard, 2019; McCafferty et al., 2018). Moreover, most current models of spike-wave initiation and maintenance neglect the potential contribution of the intralaminar nuclei to seizure initiation and maintenance despite several observations to the contrary.

In an effort to resolve structures capable of evoking spike-wave seizures, Jasper and colleagues electrically stimulated several thalamic nuclei in cats while recording EEG. By doing so in both lightly anesthetized (Jasper & Droogleever-Fortuyn, 1947) and unanesthetized (Hunter & Jasper, 1949) animals, the authors concluded that stimulation of the anterior intralaminar nuclei (i.e., central lateral, central medial and paracentral nuclei) was sufficient to evoke spike-wave seizures that outlasted the stimulus; stimulation also produced behavioral repertoires associated with absence seizures. However, stimulation of first-order thalamic nuclei did not evoke spike-wave seizures, nor did it evoke seizure-like behaviors. Consistent with these observations, lesions to the intralaminar nuclei abolish pharmacologically-induced spike-wave seizures in Sprague-Dawley rats (Banerjee & Snead, 1994); seizures persist following lesions to first-order nuclei. More recently, an EEG-fMRI study in human patients also implicates the intralaminar nuclei in the initiation of spontaneous spike-wave seizures (Tyvaert et al., 2009). Regrettably, Meeren et al. (Meeren et al., 2002) did not include intralaminar thalamic recordings during their study of spike-wave seizure propagation in the WAG/Rij rat. Nonetheless, proposing the hypothesis that the intralaminar nuclei, not cortical structures, initiate spike-wave seizures, including those occurring spontaneously (i.e., not during hyperventilation), seems premature. Indeed, the possibility that activation of cortically projecting intralaminar neurons during hyperventilation recruits cortical structures to, in turn, initiate spike-wave seizures is equally plausible (see Figure 7).

### Thalamic pH sensitivity

First-order thalamic neurons express several pH-sensitive ion channels and receptors. TASK-1 and TASK-3, two TWIK-related acid-sensitive potassium channels, with the hyperpolarization-activated cyclic nucleotide-gated (HCN) ion channel, collectively play a critical role in stabilizing the resting membrane potential of first-order thalamic neurons (Meuth et al., 2003, 2006). When activated, TASK channels hyperpolarize the membrane potential of thalamocortical neurons. In contrast, HCN channels depolarize thalamocortical neuron membrane potential. As extracellular acidification inhibits the activity of both channels, the opposing actions of TASK and HCN channels are simultaneously downregulated to yield no net effect on thalamocortical neuron membrane potential (Meuth et al., 2006), thereby stabilizing the membrane potential during acidic conditions. While not yet directly tested, the opposing actions of TASK and HCN channels also presumably stabilize thalamocortical membrane potential during alkaline conditions. Thus, while first-order thalamocortical neurons express pH-sensitive ion channels, these neurons are presumed to maintain stable membrane potentials during extracellular pH fluctuations. If true, then first-order thalamic nuclei are unlikely to support an active role in initiating hyperventilation-provoked spike-wave seizures. The extent to which higher-order thalamic nuclei express TASK and HCH channels remains unknown.

Importantly, intralaminar neurons recruited during hyperventilation-mediated alkalosis may not reflect intrinsic pH sensitivity. Instead, activation of intralaminar neurons during alkalosis may result from increased excitatory synaptic input. Intralaminar neurons receive significant, monosynaptic excitation from the midbrain reticular formation (Ropert & Steriade, 1981; Steriade & Glenn, 1982); first-order thalamic nuclei only do so negligibly (Edwards & de Olmos, 1976). Several reticular nuclei are critically important for respiration (Guyenet & Bayliss, 2015; Smith et al., 2013) and therefore provide clear rationale for testing the hypothesis that reticular-mediated excitation of the intralaminar nuclei drive hyperventilation-associated cFos expression (i.e., Figure 6). Notably, cFos expression was only observed during respiratory alkalosis (i.e., hypoxia) and not during hyperventilation associated with a normalized arterial pH (i.e., hypoxia-hypercapnia; c.f. Figures 3H and 6B). Thus, if reticular-mediated excitation of intralaminar neurons plays a role in hyperventilation-provoked spike-wave seizures, then it does so only during conditions of respiratory alkalosis. Finally, the possibility that the synaptic terminals of intralaminar-projecting afferents are pH-sensitive also warrants examination. Notably, solute carrier family transporters (SLC) shuttle H^+^ and HCO_3_^+^ across neuronal membranes and are proposed to regulate seizures, including spike-wave seizures (Cox et al., 1997; Sander et al., 2002; Sinning & Hübner, 2013). Alkaline conditions enhance excitatory synaptic transmission, an effect attributed to Slc4a8, a Na^+^-Driven Cl^-^/Bicarbonate Exchanger (Sinning et al., 2011; Sinning & Hübner, 2013), that is expressed in the presynaptic terminals of excitatory neurons, including those in the thalamus (Lein et al., 2007). Thus, the potentiation of intralaminar neuron excitation remains a plausible candidate mechanism to explain the observed cFos expression during respiratory alkalosis.

### Conclusion

In aggregate, our data support the hypothesis that spike-wave seizures are yoked to arterial pH. The observation that respiratory alkalosis activates intralaminar thalamic neurons, and that such neurons are activated by alkaline conditions, reignites a 70-year-old hypothesis wherein intralaminar neurons actively participate in the initiation and maintenance of spike-wave seizures.

## Material and Methods

### Study Design

The goal of this study was to parameterize the effect of blood gases on spike-wave seizures. To do so, we adapted a clinically observed human phenomenon in absence epilepsy patients to a rodent model of spike-wave seizures. We demonstrate that spike-wave seizure occurrence correlates with rising or falling values of PaCO_2_ and pH. Significantly, we show that neurons of the midline thalamus become activated after brief exposure to low PaCO_2_ conditions. We propose that activity among pH-sensitive neurons in the thalamus, responsive to hyperventilation-induced hypocapnia, trigger spike-wave seizures. All physiology and ECoG/EMG recordings were performed in freely behaving WAG/Rij or Wistar rats. To reduce the number of animals, rats were exposed to multiple conditions. Experimenters were blinded to the condition for all respiration and ECoG/EMG data analysis. Group and sample size were indicated in the results section.

### Animals

All procedures conformed to the National Institutes of Health *Guide for Care and Use of Laboratory Animals* and were approved by the University of Virginia Animal Care and Use Committee (Charlottesville, VA, USA). Unless otherwise stated, animals were housed at 23-25°C under an artificial 12 h light-dark cycle with food and water *ad libitum*. A colony of Wistar Albino Glaxo/from Rijswik (WAG/Rij rats) were kindly provided by Dr. Edward Bertram, University of Virginia) and maintained in the animal facilities at The University of Virginia Medical Center. Male Wistar IGS Rats were purchased from Charles River (Strain Code: #003). Plethysmography, EEG, blood gas measurements and c-Fos immunohistochemistry experiments were performed in 100+-day old WAG/Rij and Wistar rats as these ages correspond to when spike-wave seizures become robust in the WAG/Rij rat. Male and female rats were used in all experiments - no noticeable differences were observed. Of note, only male rats were used in optogenetic manipulations, as female rats were less likely to recover from surgery.

### Animal Preparation

All surgical procedures were conducted under aseptic conditions. Body temperature was maintained at 37°C. Animals were anesthetized with 1-3% isoflurane or a mixture of ketamine (75 mg/kg), xylazine (5 mg/kg) and acepromazine (1 mg/kg) administered intra-muscularly. Depth of anesthesia was monitored by lack of reflex response to a firm toe and tail pinch. Additional anesthetic was administered during surgery (25% of original dose) if warranted. All surgeries, except the arterial catheter implantation, were performed on a stereotaxic frame (David Kopf Instruments, Tujunga, CA, USA). Post-operative antibiotic (ampicillin, 125 mg/kg) and analgesia (ketoprofen, 3-5 mg/kg, subcutaneously) were administered and as needed for 3 days. Animals recovered for 1-4 weeks before experimentation.

### Electrocorticogram (ECoG) and electromyography (EMG) electrode implantation

Commercially available rat recording devices were purchased from Plastics One (Roanoke, VA, USA). Recording electrodes were fabricated by soldering insulated stainless-steel wire (A-M system, Sequim, WA, USA) to stainless-steel screws (Plastics One) and gold pins (Plastics One). On the day of surgery, a small longitudinal incision was made along the scalp. Small burr holes were drilled in the skill and ECoG recording electrodes were implanted bilaterally in the cortex. Reference electrodes were placed in the cerebellum. A twisted-looped stainless-steel wire was sutured to the superficial neck muscles for EMG recordings. The recording device was secured to the skull with dental cement and incisions were closed with absorbable sutures and/or steel clips.

### PRSX-8 lentivirus preparation

The lentivirus, *PRSX8*-hCHR2(H134R)-mCherry, was designed and prepared as described previously (Abbott et al., 2009). Lentivirus vectors were produced by the Salk Institute Viral Vector Core. The titer for the *PRSX8*-hCHR2(H134R)-mCherry lentivirus was diluted to a working concentration of 1.5 x 10^10^ TU/mL. The same batch of virus was used for all experiments included in this study.

### Virus injection and fiber optic ferrule implantation

Borosilicate glass pipettes were pulled to an external tip diameter of 25 Lim and backfilled with the lentivirus, *PRSX8*-hCHR2(H134R)-mCherry. Unilateral virus injections in the RTN were made under electrophysiological guidance of the antidromic potential of the facial nucleus (see Abbott et al., 2009; Souza et al., 2018). A total of 400 nL was delivered at three rostro caudal sites separated by 200 or 300 Lim in the RTN. Illumination of the RTN was performed by placing a 200-p.m-diameter fiber optic (Thor Labs, #BFL37-200; Newton, NJ, USA) and ferrule (Thor Labs, #CFX128-10) vertically through the cerebellum between 300-1000 Lim dorsal to RTN ChR2-expressing neurons. These animals were also implanted with ECoG/EMG recording electrodes, as detailed above. All hardware was secured to the skill with dental cement. Animals recovered for 4 weeks, as this provided sufficient time for lentivirus expression in the RTN. Virus injection location was verified post-hoc. Only animals that responded to optical stimulation, demonstrated by an increase in respiratory frequency, were included in the results.

### Physiology experiments in freely behaving rats

All experiments were performed during the dark cycle (hours 0-4) at ambient room temperature of 27°C-28°C. Rats were habituated to experimental conditions for a minimum of 4 hours, 1-2 d before experiment start. On the day of recordings, rats were briefly anesthetized with 3% isoflurane for < 5min to connect the ECoG/EMG recording head stage to a recording cable and, when necessary, to connect the fiber optic ferrule to a fiber optic cord (multimode 200p.m core, 0.39 nA) attached to a 473 nm blue laser (CrystaLaser model BC-273-060-M, Reno, NV, USA). Laser power was set to 14mW measured at the junction between the connecting fiber and the rat. Rats were then placed immediately into a whole-body plethysmography chamber (5L, EMKA Technologies, Falls Church, VA, USA). Recordings began after 1 h of habituation. The plethysmography chamber was continuously perfused with room air or protocols cycling through specific gas mixtures of O_2_, N_2_ and CO_2_ (total flow: 1.5 L/min). Mass flow controllers, operated by a custom-written Python script, regulated gas exchange. Respiratory flow was recorded with a differential pressure transducer. The respiratory signal was filtered and amplified at 0.1-100 Hz, X 500 (EMKA Technologies). Respiratory signals were digitized at 200 Hz (CED Instruments, Power1401, Cambridge, England). ECoG and EMG signals were amplified (X1000, Harvard Apparatus, Holliston, MA, USA; Model 1700 Differential Amplifier, A-M Systems), bandpass filtered (ECoG: 0.1-100 Hz; EMG: 100-300 Hz) and digitized at 200 Hz. Respiratory flow, ECoG/EMG recordings, O_2_ flow and the laser pulse protocol were captured using Spike2, 7.03 software (CED Instruments). Spike-wave seizure occurrence before and during specific conditions is shown as a peri-stimulus time histogram aligned at time = 0 at gas exchange onset or laser-on for optogenetic stimulations. Spike-wave seizure counts were quantified in 3 bins beginning +/- 15 minutes of gas exchange or laser onset. Total spike-wave seizure counts were obtained by summing the number of spike-wave seizures between −15 and 0 minutes (control) and 0 and +15 minutes (manipulation). Respiratory frequency (*f_R_*, in breaths/minute) was derived from the respiration trace. The respiration trace was divided into individual windows, each 10 seconds in duration, and a fast Fourier transform (FFT) was computed on each discrete window. The respiratory rate for each window was defined by the FFT frequency with the maximal power density. Once derived for each window, we then applied a 30-second moving average to smooth the trace. RTN neurons were optically stimulated with 10 ms pulses delivered at 20 Hz for 2 seconds, followed by 2 seconds rest. This stimulation protocol was repeated for 20 minutes.

### Femoral artery catheterization, blood gases and pH measurements

Arterial blood samples for blood gas measurements through an arterial catheter during physiological experiments. One day prior to the experiments, rats anesthetized with isoflurane (2% in pure O_2_) and a polyethylene catheter (P-10 to P-50, Clay Adams, Parsippany, NJ, USA) was introduced into the femoral artery by a small skin incision towards the abdominal aorta. The catheter was then tunneled under the skin and exteriorized between the scapulae with two inches of exposed tubing anchored with a suture. On the day of the experiment, animals were briefly anesthetized with 1-2% isoflurane to attach tubing for blood collection before placement into the plethysmography recording chamber. Arterial blood gases and pH were measured using a handheld iStat configured with CG8+ cartridges (Abbott Instruments, Lake Bluff, USA).

### cFos Histology

After exposing WAG/Rij rats to 30 minutes of hypoxia (10% O_2_; 90% N_2_) or hypoxia/hypercapnia (10% O_2_; 5% CO_2_; 75% N_2_) rats were deeply anesthetized and perfused transcardially with 4% paraformaldehyde (pH 7.4). Brains were removed and post-fixed for 12-16 h at 4 °C. 40μm horizontal sections of the thalamus (D/V depth −5.3 mm to 6.0 mm) were obtained using a Leica VT 1000S microtome (Leica Biostystems, Buffalo Grove, IL, USA) and collected in 0.1 M phosphate buffer (PB) with 0.1% sodium azide (Millipore-Sigma, St. Louis, MO, USA). Sections were then transferred to a 0.1M PB solution containing 20% sucrose for 1hr, snap-frozen and transferred to 0.1% sodium borohydride for 15 minutes. Slices were washed 2x in phosphate buffered saline (PBS). All blocking and antibody solutions were prepared in an incubation buffer of 0.1% sodium azide, 0.5% Triton X-100 and 2% normal goat serum. Sections were blocked for 4hrs at room temperature or overnight at 4°C in incubation buffer. Sections were washed 3x with PBS between primary and secondary antibody solutions. Primary antibody solutions containing rabbit anti-cFos (1:2000; Cell Signaling Technology Cat# 2250, RRID: AB_2247211, Danvers, MA, USA) and biotin (1:200, Jackson ImmunoResearch, West Grove, PA; RRID: AB_2340595) were prepared in incubation buffer and incubated overnight at 4°C. Sections were then incubated overnight in secondary antibody solutions containing donkey strepavidin-Cy3 (1:1000, Jackson ImmunoResearch; RRID: AB_2337244). Immunohistochemical controls were run in parallel on spare sections by omitting the primary antisera and/or the secondary antisera. Sections from each well were mounted and air-dried overnight. Slides were cover-slipped with VectaShield (VectorLabs, Burlingame, CA) with the addition of a DAPI counterstain. All images were captured with a Z1 Axioimager (Zeiss Microscopy, Thornwood, NY, USA) with computer-driven stage (Neurolucida, software version 10; MicroBrightfield, Inc., Colchester, VT, USA). Immunological sections were examined with a 10x objective under epifluorescence (Cy3). All sections were captured with similar exposure settings. Images were stored in TIFF format and imported into ImageJ (NIH). Images were adjusted for brightness and contrast to reflect the true rendering as much as possible.

### Calcium Imaging

pGP-AAV-syn-jGCaMP7s-WPRE (Addgene #104487-AAV9) was stereotaxically delivered to the central median thalamic nucleus in P20-30 rats with sterile microliter calibrated glass pipettes. A picospritzer (Picospritzer III, Parker Hannifin) was used to deliver 100-200 nl of virus. Three weeks later, animals were sacrificed and their brains harvested for acute brain slice preparation. Animals were deeply anesthetized with pentobarbital and then transcardially perfused with an ice-cold protective recovery solution containing the following (in mm): 92 NMDG, 26 NaHCO3, 25 glucose, 20 HEPES, 10 MgSO4, 5 Na-ascorbate, 3 Na-pyruvate, 2.5 KCl, 2 thiourea, 1.25 NaH2PO4, 0.5 CaCl2, titrated to a pH of 7.3–7.4 with HCl (Ting et al., 2014). Horizontal slices (250 μm) containing the intralaminar thalamic nuclei were cut in ice-cold protective recovery solution using a vibratome (VT1200, Leica Biosystems) and then transferred to protective recovery solution maintained at 32-34°C for 12 min. Brain slices were kept in room temperature artificial cerebrospinal fluid (ACSF) containing (in mm): 3 KCl, 140 NaCl, 10 HEPES, 10 Glucose, 2 MgCl2, 2 CaCl2. The solution was bubbled with 100% O2 and the pH was set by adding varied amounts KOH. Fluorescence signals were measured with a spinning disk confocal microscope outfitted with an sCMOS camera (ORCA-Flash4.0, Hamamatsu).

### Data analysis and statistics

Statistical analyses were performed in GraphPad Prism v7 (San Diego, CA, USA). All data were tested for normality before additional statistical testing. Statistical details, including sample size, are found in the results section and corresponding supplemental tables. Either parametric or non-parametric statistical analyses were performed. A significance level was set at 0.05. Data are expressed as mean ± SEM.

